# N^6^-methyladenosine RNA modification regulates strawberry fruit ripening in an ABA-dependent manner

**DOI:** 10.1101/2021.03.04.433875

**Authors:** Leilei Zhou, Renkun Tang, Xiaojing Li, Shiping Tian, Bingbing Li, Guozheng Qin

## Abstract

**Background:** Epigenetic marks, such as DNA methylation, play pivotal roles in regulating ripening of both climacteric and non-climacteric fruits. However, it remains unclear whether mRNA m^6^A methylation, the epitranscriptome, is functionally conserved for ripening control.

**Results:** Here we show that m^6^A methylation, which has been revealed to regulate ripening of tomato, a typical climacteric fruit, displays a dramatic change at ripening onset of strawberry, a classical non-climacteric fruit. The m^6^A modification in the coding sequence (CDS) regions appears to be ripening-specific and tends to stabilize the mRNAs, whereas m^6^A around the stop codons and within the 3’ untranslated regions is generally negatively correlated with the abundance of the mRNAs. We identified thousands of transcripts with m^6^A hypermethylation in the CDS regions, including those of *NCED5*, *ABAR*, and *AREB1* in the abscisic acid (ABA) biosynthesis and signaling pathway. We demonstrated that the methyltransferases MTA and MTB are indispensable for normal ripening of strawberry fruit, and MTA-mediated m^6^A modification promotes mRNA stability of *NCED5* and *AREB1*, while facilitates translation of *ABAR*.

**Conclusion:** Our findings uncover that m^6^A methylation regulates ripening of non-climacteric strawberry fruit by targeting ABA pathway, which is distinct from that in climacteric tomato fruit.

## Introduction

As the most prevalent chemical modification in eukaryotic messenger RNAs (mRNAs), N^6^-methyladenosine (m^6^A) has been demonstrated to functionally modulate multiple biological processes through interfering mRNA metabolism [1–4]. In mammals, m^6^A methylation has been unveiled to play critical roles in regulating various physiological and pathological processes, such as embryonic and post-embryonic development, cell circadian rhythms, and cancer stem cell proliferation [5–10]. The m^6^A marks in mammals are dominantly installed by the methyltransferase complex composed of the stable catalytic core that is formed by methyltransferase-like 3 (METTL3) and METTL14 [11–13], the Wilms tumor 1-associating protein (WTAP) [14], and other concomitant functional elements [15–17]. The removal of m^6^A marks is executed by two m^6^A demethylases, fat mass and obesity-associated protein (FTO) and alkylated DNA repair protein AlkB homolog 5 (ALKBH5) [18, 19]. The m^6^A modification are recognized by the reader proteins, such as YTH-domain family proteins and specific RNA binding proteins (RBPs), which mediate the downstream effects of m^6^A methylation [4]. In plants, the m^6^A methylation machineries have been characterized in the model plant *Arabidopsis thaliana* to modulate development processes such as shoot stem cell proliferation, trichome branching, and floral transition [20–23]. Moreover, m^6^A has been demonstrated to play pivotal roles in mediating sporogenesis in rice [24] and regulating stress responses in maize [25]. By contrast, the m^6^A methylation machineries as well as the characteristics and functions of m^6^A in regulating physiological processes of horticultural crops remain largely unknown.

Fleshy fruits, which are enriched with nutrients, such as flavor compounds, fiber, vitamins and antioxidants, represent one of the commercially valuable structures of horticultural crops. As an important component of diets, fleshy fruits are indispensable for human health [26]. The ripening of fleshy fruits, which is characterized by dramatic changes in color, texture, flavor and aroma compounds [27], is a complex, genetically programmed process that impacts fruit nutritional quality and shelf life. Fruit ripening is regulated by both environmental and internal cues, including light, temperature, phytohormones, and developmental genes [28, 29]. Based on the different ripening mechanisms, fruits are classified into two groups: climacteric (e.g. tomato, apple, banana, and avocado) and non-climacteric (e.g. strawberry, grape, and citrus) [30]. Phytohormone ethylene is essential for the ripening of climacteric fruits [31, 32], and substantial insights have been made toward ethylene biosynthesis, ethylene perception and signal transduction, and downstream gene regulation [33]. In comparison, the ripening of non-climacteric fruits is thought to be abscisic acid (ABA)-dependent [32, 34], although the regulation of ABA pathway is poorly understood. A comprehensive understanding of the common regulatory mechanisms underlying ripening in climacteric and non-climacteric species has great potential for improving fruit quality and maintaining shelf-life.

Recently, it was shown that epigenetic marks, including DNA methylation and histone posttranslational modifications, play critical roles in the regulation of fruit ripening [35]. In a previous study, we uncovered that mRNA m^6^A methylation, which is considered as an mRNA “epitranscriptome”, exhibits dynamic changes during fruit ripening of tomato, a typical climacteric fruit [36]. Mutation of *SlALKBH2*, the m^6^A RNA demethylase gene, delays fruit ripening [36], indicating that m^6^A modification participates in the ripening control of climacteric fruit tomato. However, whether m^6^A is evolutionarily conserved among different types of fruits has not been defined. Moreover, the regulatory function of m^6^A in ripening of non-climacteric fruits remains elusive.

In the present study, we performed transcriptome-wide m^6^A methylation (m^6^A methylome) in strawberry, a classical non-climacteric fruit, and revealed that m^6^A represents a prevalent modification in the mRNAs of strawberry fruit. Compared to the dynamic changes in m^6^A modification around the stop codons or within the 3’ untranslated regions during the ripening of tomato fruit, a specific enrichment of m^6^A in the coding sequence (CDS) region, which tends to be positively correlated with the abundance of the transcripts, was observed in the ripe strawberry fruit. We demonstrated that, mediated by the methyltransferases MTA and MTB, m^6^A modification enhances mRNA stability or promotes translation efficiency of genes in the ABA biosynthesis and signaling pathway, thereby facilitating the ripening of strawberry fruit. Our study uncovers the regulatory effects of m^6^A methylation on non-climacteric strawberry fruit ripening, and reveals a direct role for m^6^A methylation in the regulation of key elements in the ABA pathway.

## Results

### m^6^A methylation is a common feature of mRNAs in strawberry fruit

To investigate whether m^6^A methylation participates in modulating ripening of non-climacteric fruits, we performed m^6^A-seq [37] to characterize m^6^A methylomes on diploid woodland strawberry (*Fragaria vesca*) at three developmental stages, i.e. S6 (the growth stage 6, approximately 15 days post-anthesis (DPA)), RS1 (the ripening stage 1, 21 DPA), and RS3 (the ripening stage 3, 27 DPA) (Fig. 1a) [38]. The transition from S6 to RS1 represents the initiation of ripening, and that from RS1 to RS3 indicates the phase after ripening initiation. The m^6^A methylome libraries were prepared with three independently biological replicates and subjected to massively parallel sequencing according to the standard m^6^A-seq protocols [37]. High Pearson correlation coefficients were observed between biological replicates, indicating reliable repeatability (Additional file 1: Figure S1, S2). A total of 24-37 million raw reads were generated for each library (Additional file 2: Table S1), and this sequencing depth is comparative to that observed in mammal (11-24 million reads) [39], rice (23-47 million reads) [40], and tomato (20-30 million reads) [36]. After adaptor trimming and reads filtration, 24-37 million clean reads were remained at each library, and almost 95% of these reads were uniquely aligned to the strawberry genome v1.1, representing high mapping quality (Additional file 2: Table S1). The peak-calling algorithm was used to identify m^6^A peaks with an estimated false discovery rate (FDR) < 0.05 [37], and only those consistently detected in all three biological replicates, which we called confident m^6^A peaks, were used for subsequent analysis. We identified 9778, 10853 and 10095 confident m^6^A peaks within 8934, 8990 and 8374 gene transcripts, in fruit at S6, RS1, and RS3, respectively (Fig. 1b; Additional file 2: Table S2-S4).

**Figure 1.**
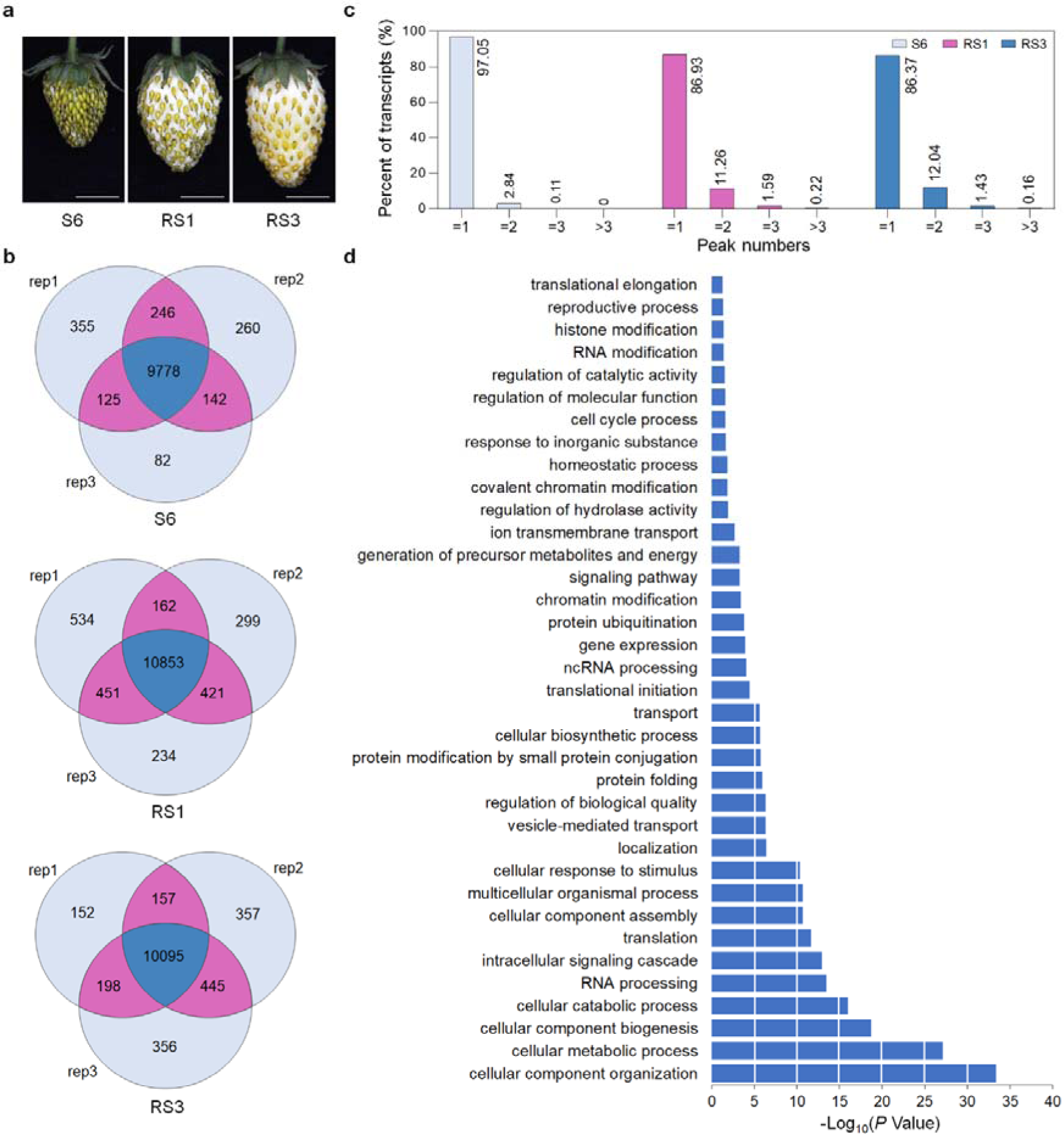
Transcriptome-wide m^6^A methylomes in strawberry fruit. **a** The representative photographs of fruit at different developmental stages. S6, the growth stage 6; RS1, the ripening stage 1; RS3, the ripening stage 3. Scale bar = 1 cm. **b** Venn diagrams depicting the overlap of m^6^A peaks from three independent m^6^A-seq experiments on fruit at the three developmental stages. Rep, replicate. **c** Percentage of the m^6^A-containing transcripts containing various m^6^A peak numbers among samples. **d** Gene Ontology (GO) enrichment for the m^6^A-modified transcripts identified in m^6^A-seq.

The m^6^A-seq results were validated by independent m^6^A immunoprecipitation followed by qPCR (m^6^A-IP-qPCR) analysis. Three m^6^A peak-containing transcripts, as well as three m^6^A peak-free transcripts were randomly selected and examined (Additional file 1: Figure S3a). As expected, m^6^A enrichment was only observed in transcripts containing m^6^A peaks, but not in those without m^6^A peaks (Additional file 1: Figure S3b), indicating that our m^6^A-seq data were accurate and robust.

Based on the parallel RNA-seq analyses, we estimated that the transcriptome of diploid woodland strawberry contains 0.6-0.7 m^6^A peaks per actively expressed transcript, which shows FPKM (fragments per kilobase of transcript per million fragments mapped) ≥ 1 (Additional file 2: Table S5). These levels are comparable with those observed in *Arabidopsis* or tomato [36, 41], demonstrating that m^6^A modification is a common feature of mRNA in strawberry fruit. Most of the m^6^A-containing transcripts (>86%) possess one m^6^A peak. Intriguingly, the percentage of transcripts harboring two or more m^6^A peaks increases dramatically when the strawberry fruit turn to ripen, which changes from 2.95% at S6 to 13.07% at RS1, and 13.63% at RS3 (Fig. 1c), raising the possibility that new m^6^A peaks generate at the initiation stage of ripening. Gene Ontology (GO) enrichment analysis indicated that m^6^A modification appears in genes in multiple signaling pathways and cellular processes (Fig. 1d).

### m^6^A distribution exhibits a dramatic change at the initiation stage of strawberry fruit ripening

We then evaluated the distribution of m^6^A peaks in the whole transcriptome of strawberry fruit. The transcript was divided into five non-overlapping segments: transcription start site (TSS, 100-nucleotide window centered on the TSS), 5’ untranslated region (UTR), coding sequence (CDS), stop codon (100-nucleotide window centered on the stop codon), and 3’ UTR. As shown in Fig. 2a, m^6^A modifications in all three samples (fruit at S6, RS1, and RS3 stages) were highly enriched around the stop codon and within the 3’ UTR, but the percentage of peak summits within these regions declined markedly in the ripening process (from S6 to RS1 or RS3; indicated by green arrowheads). Surprisingly, a substantial increase in percentage of peak summits was concurrently observed in the CDS region adjacent to the start codon in fruit at RS1 or RS3 compared to those at S6 (indicated by red arrowheads). This is different from that observed in tomato, which shows no dramatic changes in percentage of peak summits at the initiation stage of ripening [36].

**Figure 2.**
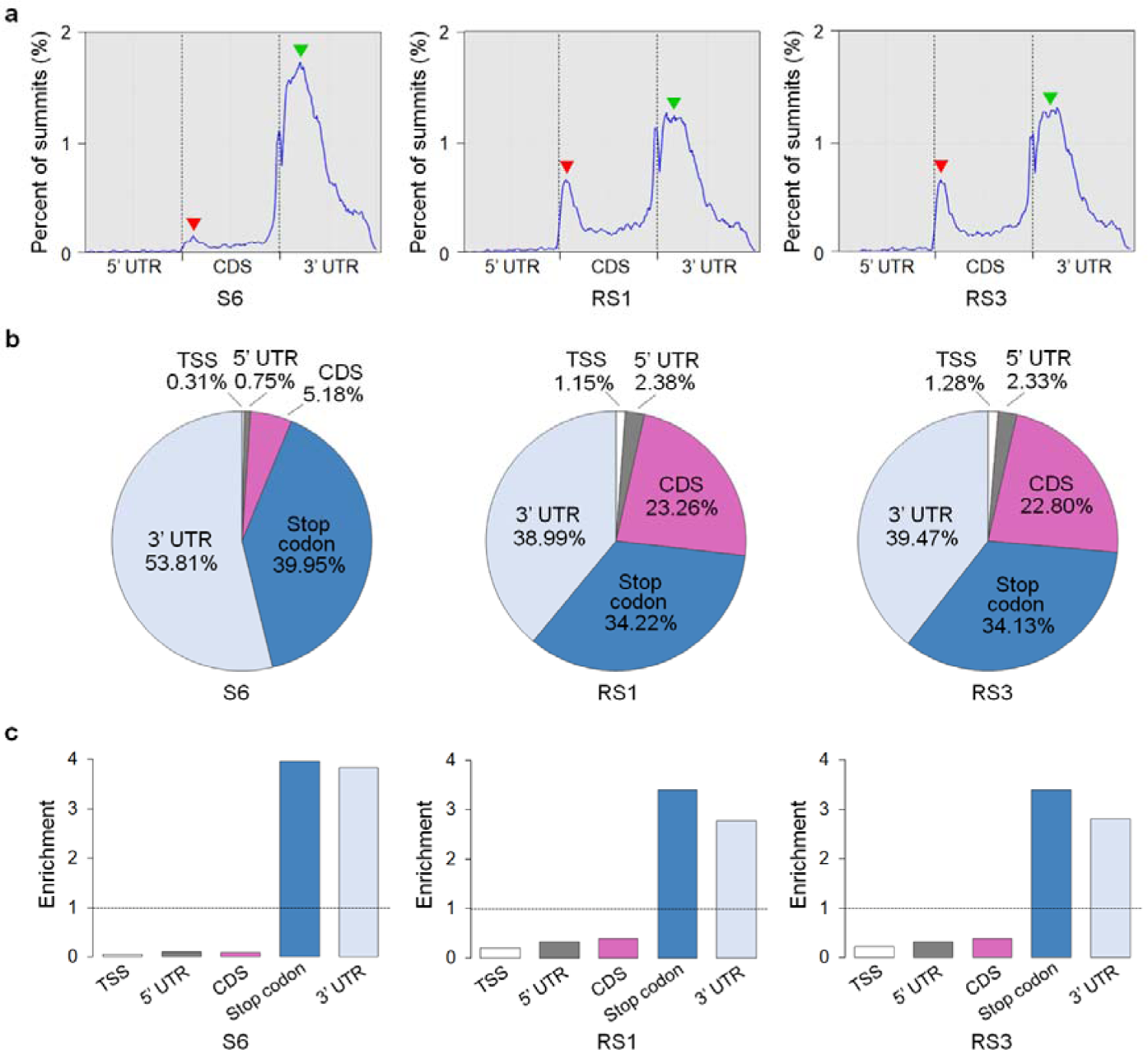
Dynamic distribution of m^6^A during strawberry fruit ripening. **a** Metagenomic profiles of m^6^A peak summit distribution along transcripts. UTR, untranslated region; CDS, coding sequence. The red arrows indicate the changes in m^6^A distribution in the CDS region adjacent to the start codon in the ripening process, while the green arrows show the changes in m^6^A distribution around the stop codon or within the 3’ UTR. S6, the growth stage 6; RS1, the ripening stage 1; RS3, the ripening stage 3. **b** and **c** The percentage (**b**) and relative enrichment (**c**) of m^6^A peak summits in five non-overlapping transcript segments. TSS, transcription start site.

The percentage of peak summits locating in the CDS region increased from 5.18% to 23.26% from S6 to RS1, but displayed no distinct changes from RS1 to RS3 (Fig. 2b). In comparison, the percentage of peak summits felling into the stop codon region and the 3’ UTR decreased from 39.95% and 53.81%, to 34.22% and 38.99%, respectively, from S6 to RS1, with no obvious changes observed from RS1 to RS3 (Fig. 2b). After segment normalization by the relative fraction that each segment occupied in the transcriptome, the m^6^A enrichment in fruit at RS1 or RS3 consistently exhibit a preferential localization in CDS region, besides enriched around the stop codon and within the 3’ UTR (Fig. 2c). Thus, the transcriptome-wide m^6^A distribution displays a dramatic change at the initiation stage of ripening, but not after ripening initiation.

Several high-confidence sequence motifs were identified within the m^6^A peaks (Additional file 1: Figure S4), by using the hypergeometric optimization of motif enrichment (HOMER; http://homer.ucsd.edu/homer/) [42]. The conserved RR<u>A</u>CH consensus sequence observed in mammals [43], where R represents adenosine (A) or guanosine (G), underlined A indicates m^6^A, and H represents A, cytidine (C), or uridine (U), appears in the list, whereas the UGUA sequence motif found in *Arabidopsis* [22], tomato [36], and maize [25] was not identified.

These data suggest the complexity of m^6^A modification among various species.

### m^6^A methylation overall affects mRNA abundance during the ripening of strawberry fruit

To gain insight into the potential roles of m^6^A in the regulation of strawberry fruit ripening, we next sought for transcripts showing differential m^6^A peaks, which exhibit a fold change ≥ 1.5 and *P* value < 0.05 in m^6^A enrichment between the samples, by comparing the m^6^A methylomes. A total of 1608 hypermethylated m^6^A peaks and 865 hypomethylated m^6^A peaks, corresponding to 1398 and 790 transcripts, respectively, were identified in fruit at RS1 compared to those at S6 (Fig. 3a; Additional file 2: Table S6, S7). By contrast, only 113 hypermethylated m^6^A peaks and 102 hypomethylated m^6^A peaks, which were distributed in 107 and 90 transcripts, respectively, were identified in fruit at RS3 compared to those at RS1 (Fig. 3b; Additional file 2: Table S8, S9). These results confirmed that substantial changes in overall m^6^A methylome occurred at the initiation stage of ripening (from S6 to RS1), but not after ripening initiation (from RS1 to RS3).

**Figure 3.**
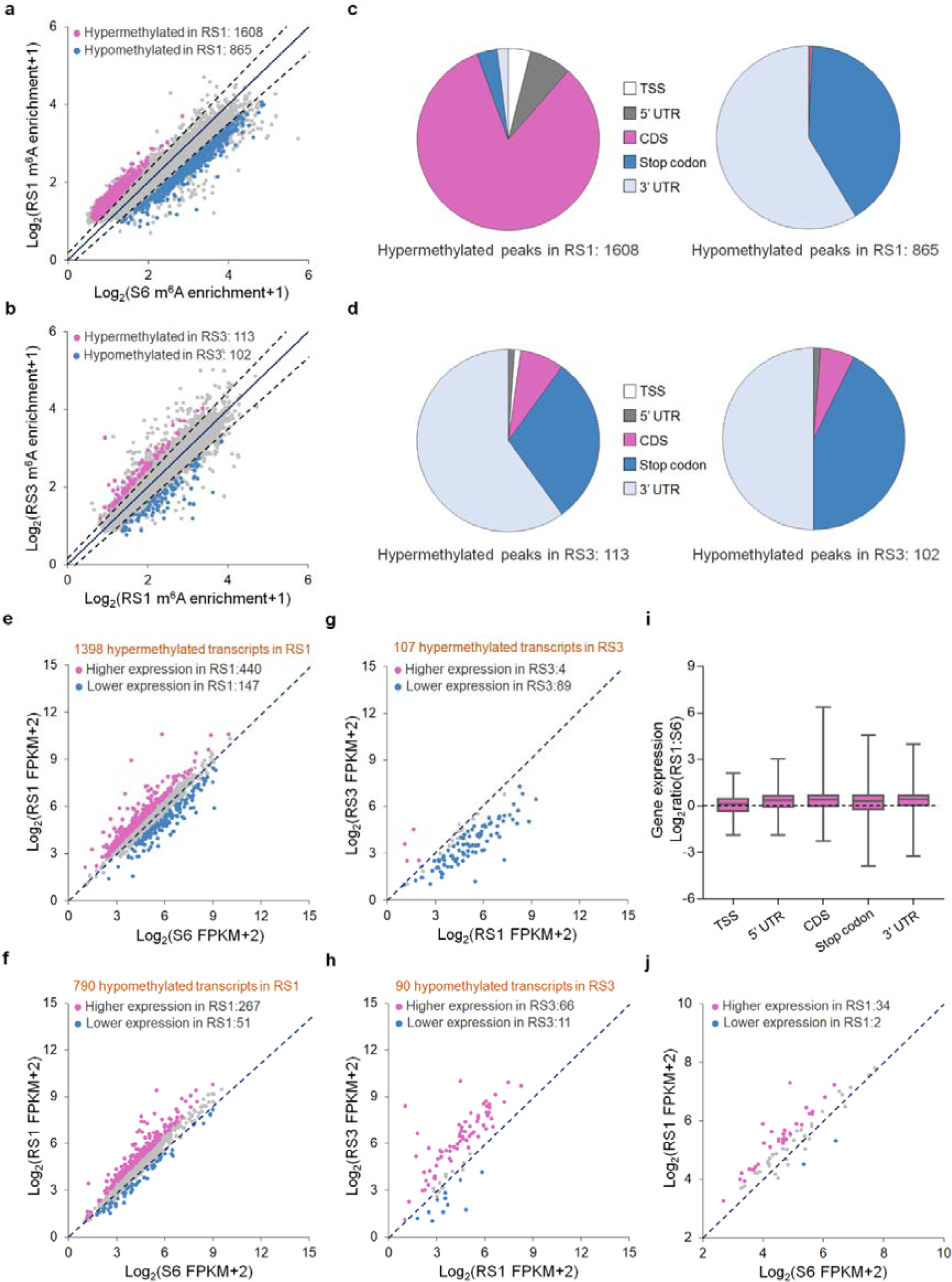
Correlation between m^6^A modification and mRNA abundance in strawberry fruit. **a** Scatter plots showing hypermethylated (red) and hypomethylated (blue) m^6^A peaks in fruit at RS1 stage compared to those at S6 stage. S6, the growth stage 6; RS1, the ripening stage 1. **b** Scatter plots depicting hypermethylated (red) and hypomethylated (blue) m^6^A peaks in fruit at RS3 stage compared to those at RS1 stage. RS3, the ripening stage 3. **c** and **d** The distribution characteristics of the differential m^6^A peaks shown in **a** (**c**) and **b** (**d**). TSS, transcription start site; UTR, untranslated region; CDS, coding sequence. **e** and **f** Scatter plots displaying the expression of transcripts containing hypermethylated (**e**) or hypomethylated (**f**) m^6^A peaks shown in **a**. Transcripts with significantly higher and lower levels (fold change ≥ 1.5; *P* value < 0.05) in fruit at RS1 stage compared to those at S6 stage are highlighted in red and blue, respectively. **g** and **h** Scatter plots showing the expression of transcripts harboring hypermethylated (**g**) or hypomethylated (**h**) m^6^A peaks shown in **b**. Transcripts with significantly higher and lower levels (fold change ≥ 1.5; *P* value < 0.05) in fruit at RS3 stage compared to those at RS1 stage are highlighted in red and blue, respectively. **i** Gene expression ratios based on the distributions of differential m^6^A peaks between fruit at RS1 and S6 stages. **j** Scatter plots exhibiting the expression of transcripts concurrently containing hypermethylated m^6^A peaks in the CDS region and hypomethylated m^6^A peaks around the stop codon or within the 3’ UTR in fruit at RS1 stage compared to those at S6 stage. Transcripts with significantly higher and lower levels (fold change ≥ 1.5; *P* value < 0.05) in fruit at RS1 stage compared to those at S6 stage are highlighted in red and blue, respectively. Gene expression analysis in **e**-**j** was performed by RNA-seq. FPKM, fragments per kilobase of exon per million mapped fragments.

The 1608 hypermethylated m^6^A peaks were highly enriched (83.04%) in the CDS region, whereas the 865 hypomethylated m^6^A peaks were mainly distributed around the stop codon (40.66%) or within the 3’ UTR (58.59%) (Fig. 3c). This is in accordance with the results of m^6^A distribution (Fig. 2), showing that the percentage of peak summits locating in the CDS region increased sharply, while that around the stop codon or within the 3’ UTR declined, during from S6 to RS1. Interestingly, of the 1608 hypermethylated m^6^A peaks, 1424 (88.56%) fell into the newly generated peaks, which represents ripening-specific peaks (Additional file 2: Table S10). For the differential m^6^A peaks identified after ripening initiation, both the hypermethylated and hypomethylated m^6^A peaks were highly enriched (over 90%) around the stop codon or within the 3’ UTR (Fig. 3d).

Accumulating evidences have confirmed that m^6^A deposition affects mRNA abundance [1, 22, 41, 44]. To assess the potential correlation between m^6^A modification and mRNA abundance in strawberry fruit, we compared the list of transcripts harboring altered m^6^A methylation with the differentially expressed genes (fold change ≥ 1.5 and *P* value < 0.05) obtained from our parallel RNA-seq analyses (Additional file 2: Table S11, S12). Among the 1398 transcripts with hypermethylated m^6^A peaks in fruit at RS1 compared to those at S6, 440 and 147 transcripts displayed higher and lower expression levels, respectively (Fig. 3e; Additional file 2: Table S13). The distribution features of m^6^A modifications in these transcripts (Fig. 3c) suggest that m^6^A depositions in CDS region overall exhibit a positive effect on mRNA abundance. Accordingly, among the 790 transcripts showing hypomethylated m^6^A peaks in fruit at RS1 compared to those at S6, 267 transcripts showed higher expression levels, whereas only 51 transcripts exhibited lower expression levels (Fig. 3f; Additional file 2: Table S14). The negative role of m^6^A modifications on mRNA abundance was also observed in transcripts with hypermethylated (Fig. 3g) or hypomethylated (Fig. 3h) m^6^A peaks that were identified after ripening initiation (Additional file 2: Table S15, S16). Considering the distribution characteristics of m^6^A within these transcripts (Fig. 3c, d), it was proposed that, consistent with the observation in tomato fruit [36], m^6^A depositions around the stop codon or within the 3’ UTR are generally negatively correlated with the abundance of the transcripts in strawberry fruit.

To further evaluate the potential correlation between m^6^A deposition and mRNA abundance, the changes in transcript levels were separately explored, based on the m^6^A distributions, for transcripts showing differential m^6^A modification at the initiation stage of ripening (from S6 to RS1). Gene expression profiles showed that most of the differential m^6^A-modified transcripts (approximately 75%) of the 5’ UTR, CDS, stop codon, and 3’ UTR categories exhibited relatively higher transcript levels (Fig. 3i), suggesting that m^6^A methylation overall affects mRNA abundance upon initiation of strawberry fruit ripening. Notably, 73 transcripts concurrently harboring hypermethylated m^6^A peaks in the CDS region and hypomethylated m^6^A peaks around the stop codon or within the 3’ UTR were identified at this process. Of these, 34 transcripts were expressed at higher levels in fruit at RS1 compared to those at S6, while only 2 were expressed at lower levels (Fig. 3j), implying that the effects of m^6^A methylation on the abundance of the transcripts may be overlaid.

Notably, hundreds of ripening-induced and ripening-repressed genes, which display significantly higher or lower expression in RS1 compared to S6 (Additional file 2: Table S11), exhibit differential m^6^A modification (Additional file 2: Table S13, S14), implicating the involvement of m^6^A methylation in the regulation of strawberry fruit ripening.

### Genes in ABA biosynthesis and signaling pathway exhibit differential m^6^a methylation upon ripening initiation

The plant hormone ABA has been elucidated to be essential for strawberry fruit ripening [32]. Two core ABA signal transduction pathways, the ‘ABA-PYR/PYL-PP2C-SnRK2-AREB/ABF’ pathway [45, 46] and the ‘ABA-ABAR-WRKY40-ABI5’ pathway [47], have been proposed in *Arabidopsis*, respectively (Fig. 4a). In the m^6^A-seq analysis, we found that transcripts of key genes in ABA biosynthesis and signaling pathway, including *9-cis-epoxycarotenoid dioxygenase 5* (*NCED5*) [38], *putative ABA receptor* (*ABAR*) [34], and *ABA-responsive element-binding protein 1* (*AREB1*) [46], exhibit hypermethylation in the CDS region at the initiation stage of strawberry fruit ripening (from S6 to RS1) (Fig. 4b, c). *NCED5* encodes the rate-limiting enzyme for ABA biosynthesis, while *ABAR* and *AREB1* encode an ABA receptor and an ABA-responsive element, respectively. All three genes have been proposed to regulate strawberry fruit ripening [34, 38, 48]. The differential m^6^A modifications were confirmed by m^6^A-IP-qPCR analysis (Fig. 4d). The transcript levels of *NCED5* and *AREB1*, but not *ABAR*, increased significantly in fruit at RS1 compared to those at S6 as revealed by both RNA-seq (Fig. 4e) and quantitative RT-PCR analyses (Fig. 4f), indicating a positive correlation between m^6^A depositions and mRNA abundances. It is noteworthy that similar results of m^6^A enrichment as well as transcript levels of these three genes were found upon ripening initiation of the octoploid cultivated strawberry (*Fragaria × ananassa*) (Additional file 1: Figure S5).

**Figure 4.**
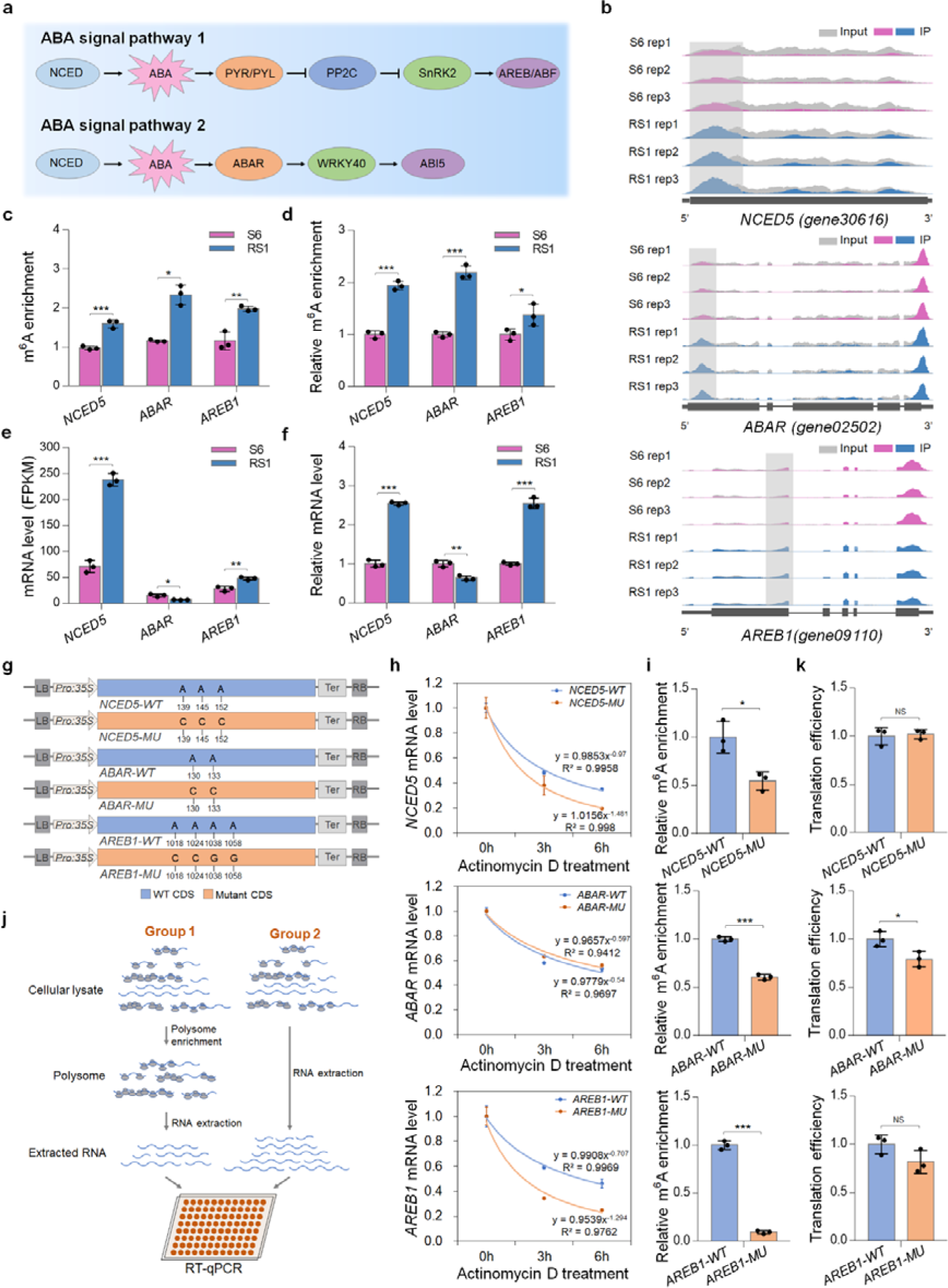
m^6^A modification facilitates mRNA stability or translation of genes in ABA pathway. **a** A brief model of the two core ABA signaling pathways in plants. **b** Integrated Genome Browser (IGB) tracks showing the distribution of m^6^A reads in transcripts of the *9-cis-epoxycarotenoid dioxygenase 5* (*NCED5*), *putative ABA receptor* (*ABAR*), and *ABA-responsive element-binding protein 1* (*AREB1*). The hypermethylated m^6^A peaks (fold change ≥ 1.5; *P* value < 0.05) in fruit at RS1 stage compared to those at S6 stage are indicated by shadow boxes. S6, the growth stage 6; RS1, the ripening stage 1. Rep, replicate. **c** m^6^A enrichment for *NCED5*, *ABAR*, and *AREB1* from m^6^A-seq data. **d** Validations of the m^6^A enrichment by m^6^A-immunoprecipitation (IP)-qPCR. **e** and **f** Transcript levels of *NCED5*, *ABAR*, and *AREB1* determined by RNA-seq (**e**) and quantitative RT-PCR (**f**). FPKM, fragments per kilobase of exon per million mapped fragments. The *ACTIN* gene served as an internal control in f. **g** Schematic diagram of the expression cassettes used for mRNA stability assay. The intact (WT) or mutated (MU) cDNA fragments of *NCED5*, *ABAR*, and *AREB1* were separately cloned into the pCambia2300 vector driven by the CaMV 35S promoter. The potential m^6^A sites identified in m^6^A-seq were mutated from adenosine (A) to cytidine (C) or guanine (G) using site-directed mutagenesis kit. **h** Determination of mRNA stability for *NCED5*, *ABAR*, and *AREB1*. The intact (WT) or mutated (MU) cDNA fragments were expressed in *Nicotiana benthamiana* leaves. After actinomycin D treatment at an indicated time point, the total RNAs were extracted and submitted to quantitative RT-PCR assay with the *N. benthamiana ACTIN* gene serving as an internal control. **i** m^6^A-IP-qPCR assay showing the relative m^6^A enrichment in the intact or mutated transcripts. **j** Brief workflow for analysis of translation efficiency. **k** Translation efficiency of *NCED5*, *ABAR*, and *AREB1*. Translation efficiency was expressed as the abundance ratio of mRNA in the polysomal RNA versus the total RNA. Data are presented as mean ± standard deviation (n = 3). Asterisks indicate significant differences (\**P* < 0.05, \*\**P* < 0.01, \*\*\**P* < 0.001; Student’s t test).

To better understand how m^6^A methylation affects the abundance of the transcripts, we examined the mRNA stability of *NCED5*, *AREB1*, and *ABAR* by monitoring the degradation rate of mRNAs in the presence of transcription inhibitor actinomycin D. The intact CDS sequences of the three genes and their mutated forms in which the potential m^6^A sites identified in m^6^A-seq were mutated from A to cytidine (C) or guanosine (G) were separately inserted into the vector for transient expression in the *Nicotiana benthamiana* leaves (Fig. 4g).

As shown in Fig. 4h, the mRNAs of *NCED5*, *AREB1*, and *ABAR* degraded obviously after actinomycin D treatment. The degradation rate was substantially increased for mRNAs of *NCED5* and *AREB1*, but not *ABAR*, in the mutated form (Fig. 4h), concomitant with significantly diminished m^6^A depositions (Fig. 4i), indicating that site-specific m^6^A modification stabilizes mRNAs of *NCED5* and *AREB1*.

The m^6^A modification in the CDS region has been demonstrated to affect translation efficiency beyond mRNA stability [3]. We next investigated whether m^6^A methylation modulates translation efficiency of *NCED5*, *AREB1*, and *ABAR*, which was determined by calculating the abundance ratio of mRNA in the polysomal RNA versus the total RNA [49] (Fig. 4j). The translation efficiency of *ABAR* rather than *NCED5* and *AREB1* was significantly decreased when the potential m^6^A modification site was mutated (Fig. 4k), demonstrating that m^6^A methylation facilitates translation of *ABAR*. Collectively, these data suggest that critical genes in ABA pathway undergo m^6^A-mediated post-transcriptional regulation, which promotes mRNA stability or facilitates translation.

### Characterizations of m^6^A methyltransferases in strawberry fruit

Having observed the changes in m^6^A methylation in a large number of transcripts (Fig. 3), including those of ABA biosynthesis and signaling genes (Fig. 4), during the ripening of strawberry fruit, we next explored the mechanistic basis. We speculate that the ripening-specific hypermethylation in the CDS region is regulated by mRNA m^6^A methyltransferases. In mammals, METTL3 and METTL14 form a stable heterodimer wherein METTL3 serves as the m^6^A catalytic subunit and METTL14 facilitates RNA binding [12, 13] (Fig. 5a). The m^6^A methyltransferase complexes are conserved between mammals and plants [50], and the *Arabidopsis* mRNA adenosine methyltransferase (MTA) and MTB appears to be the homologs of METTL3 and METTL14, respectively [51, 52]. We identified the single homologs of *Arabidopsis* MTA and MTB in the genome of the diploid woodland strawberry. Phylogenetic analysis indicated that MTA and MTB exhibit high similarity among plant species and are evolutionarily conserved with mammal METTL3 and METTL14 (Fig. 5b). Both MTA and MTB in strawberry contain the highly conserved MT-A70 domain (Additional file 1: Figure S6), which displays extremely similar protein sequences to those observed in mammals and *Arabidopsis* (Additional file 1: Figure S7).

**Figure 5.**
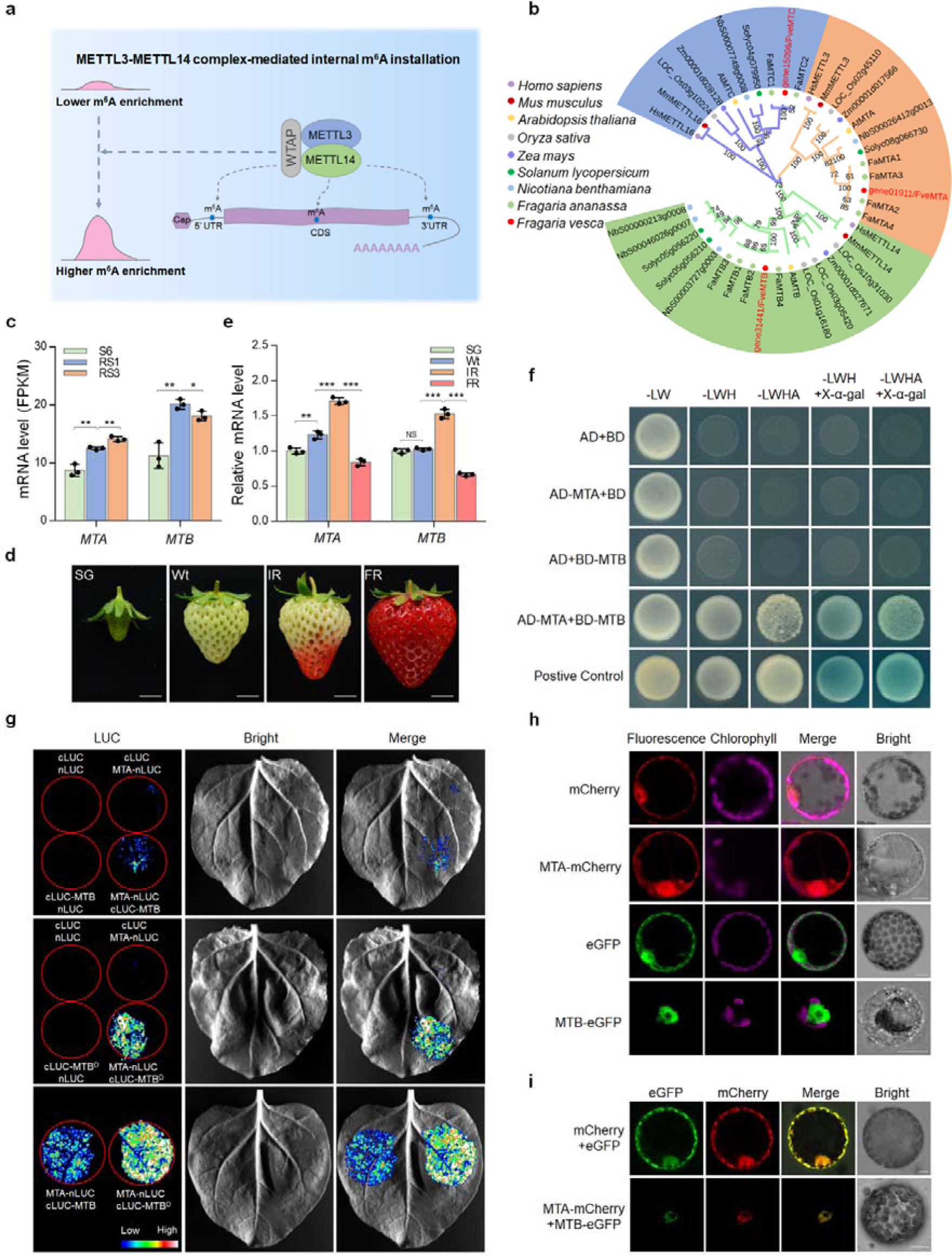
The m^6^A methyltransferase MTA interacts with MTB in strawberry. **a** The working model for m^6^A installations mediated by the methyltransferase complex in mammals. The m^6^A methyltransferases METTL3 and METTL14 interact and function as the stable catalytic core for internal m^6^A installations in the form of heterodimer. **b** Phylogenetic analysis of eukaryotic m^6^A methyltransferases. The phylogenetic tree was generated by MEGA (version 5.2). Bootstrap values from 1000 replications for each branch are presented. Species names are abbreviated as follows: Hs, *Homo sapiens*; Ms, *Mus musculus*; At, *Arabidopsis thaliana*; Os, *Oryza sativa*; Zm, *Zea mays*; Sl, *Solanum lycopersicum*; Nb, *Nicotiana benthamiana*; Fa, *Fragaria × ananassa*; Fve, *Fragaria vesca*. **c** Transcript levels of the m^6^A methyltransferase genes *MTA* and *MTB* in diploid woodland strawberry at different developmental stages revealed by RNA-seq. S6, the growth stage 6; RS1, the ripening stage 1; RS3, the ripening stage 3. **d** Representative images of octoploid cultivated strawberry fruit at various developmental stages. SG, small green; Wt, white; IR, initial red; FR, full red. Scale bar = 1 cm. **e** Transcript levels of *MTA* and *MTB* in octoploid strawberry fruit determined by quantitative RT-PCR. The *ACTIN* gene was used as an internal control. Data are presented as mean ± standard deviation (n = 3). Asterisks indicate significant differences (\**P* < 0.05, \*\**P* < 0.01, \*\*\**P* < 0.001; Student’s t test). **f** Y2H assay revealing the interactions between MTA and MTB. The MTA fused with the activation domain (AD) of GAL4 (AD-MTA) and the MTB fused with the binding domain (BD) of GAL4 (BD-MTB) were co-expressed in yeast. The transformants were grew on SD/-Leu/-Trp (-LW), and further selected on SD/-Leu/-Trp/-His (-LWH) and SD/-Leu/-Trp/-His/-Ade (-LWHA) with or without X-α-gal. **g** LCI assay revealing the interactions between MTA and MTB. The MTA fused with the N-terminus of LUC (MTA-nLUC) was co-expressed with the MTB or its MT-A70 domain fused with the C-terminus of LUC (cLUC-MTB or cLUC-MTB^D^) in *N.benthamiana* leaves. **h** and **i** Subcellular localization (**h**) and colocalization (**i**) of MTA and MTB. The MTA-mCherry or/and MTB-eGFP fusion proteins were transiently expressed into *N. benthamiana* leaves. The *N. benthamiana* leaves expressing eGFP or/and mCherry were used as the negative control. Scale bar = 10 µm.

Transcriptome analysis showed that *MTA* and *MTB* increased significantly at the initiation stage of ripening (from S6 to RS1) (Fig. 5c). The homolog genes of *MTA* and *MTB* in the octoploid cultivated strawberry also exhibited increased expression from white (Wt) stage to initial red (IR) stage (Fig. 5d, e), suggesting that the two methyltransferases may play important roles in modulating strawberry fruit ripening.

To explore the possibility that MTA and MTB in strawberry function in the form of heterodimer as METTL3 and METTL14 in mammals [12, 13], we subsequently analyzed the interactions between MTA and MTB using the yeast two-hybrid (Y2H) system. As shown in Fig. 5f, the yeast cells co-expressing AD-MTA and BD-MTB, but not the negative controls, displayed normal growth on the selective SD/-Leu-Trp-His (-LWH) and SD/-Leu-Trp-His-Ade (-LWHA) solid medium and turn to blue with the addition of X-α-gal, indicating that MTA interacts with MTB. The interactions between MTA and MTB were further verified by the split luciferase complementation imaging (LCI) assay, in which the luciferase activity was detected when MTA-nLUC and cLUC-MTB were co-expressed in *N. benthamiana* leaves (Fig. 5g). It should be noted that, compared with the MT-A70 domain of MTB, the full-length MTB protein exhibit relatively weaker combining capacity with MTA (Fig. 5g). Subcellular localization analysis by transiently expressing MTA-mCherry and MTB-eGFP fusion proteins in *N. benthamiana* leaves showed that MTA is present in both the nucleus and cytoplasm, while the MTB protein is specifically localized in the nucleus (Fig. 5h). Interestingly, when MTA-mCherry was co-expressed with MTB-eGFP, the two proteins tend to colocalization in the nucleus (Fig. 5i).

### m^6^A methyltransferases positively regulate strawberry fruit ripening

We subsequently examined the function of MTA and MTB in strawberry fruit ripening. The RNA interference (RNAi) and overexpression constructs of *MTA* or *MTB* under the control of a 35S cauliflower mosaic virus promoter were agroinfiltrated into the octoploid strawberry fruit according to the reported procedures [53]. By comparing the fruits of RNAi with the control, we found that suppression of either *MTA* or *MTB* delayed fruit ripening (Fig. 6a). A visible color change was observed at the seventh days after agroinfiltration in the control, whereas the *MTA* or *MTB* RNAi fruits were almost green at this stage (Fig. 6a). Conversely, overexpression of *MTA* or *MTB* accelerated fruit ripening (Fig. 6a). Gene expression analysis indicated that *MTA* and *MTB* were successfully silenced in the RNAi fruits while enhanced in the overexpressed fruits (Fig. 6b). The ripening genes *chalcone synthase* (*CHS*) and *polygalacturonase 1* (*PG1*) displayed a noticeable decrease in the *MTA* or *MTB* RNAi fruits, but were dramatically enhanced in the overexpressed fruits (Fig. 6b). These results suggest that *MTA* and *MTB* are necessary for normal fruit ripening of strawberry.

**Figure 6.**
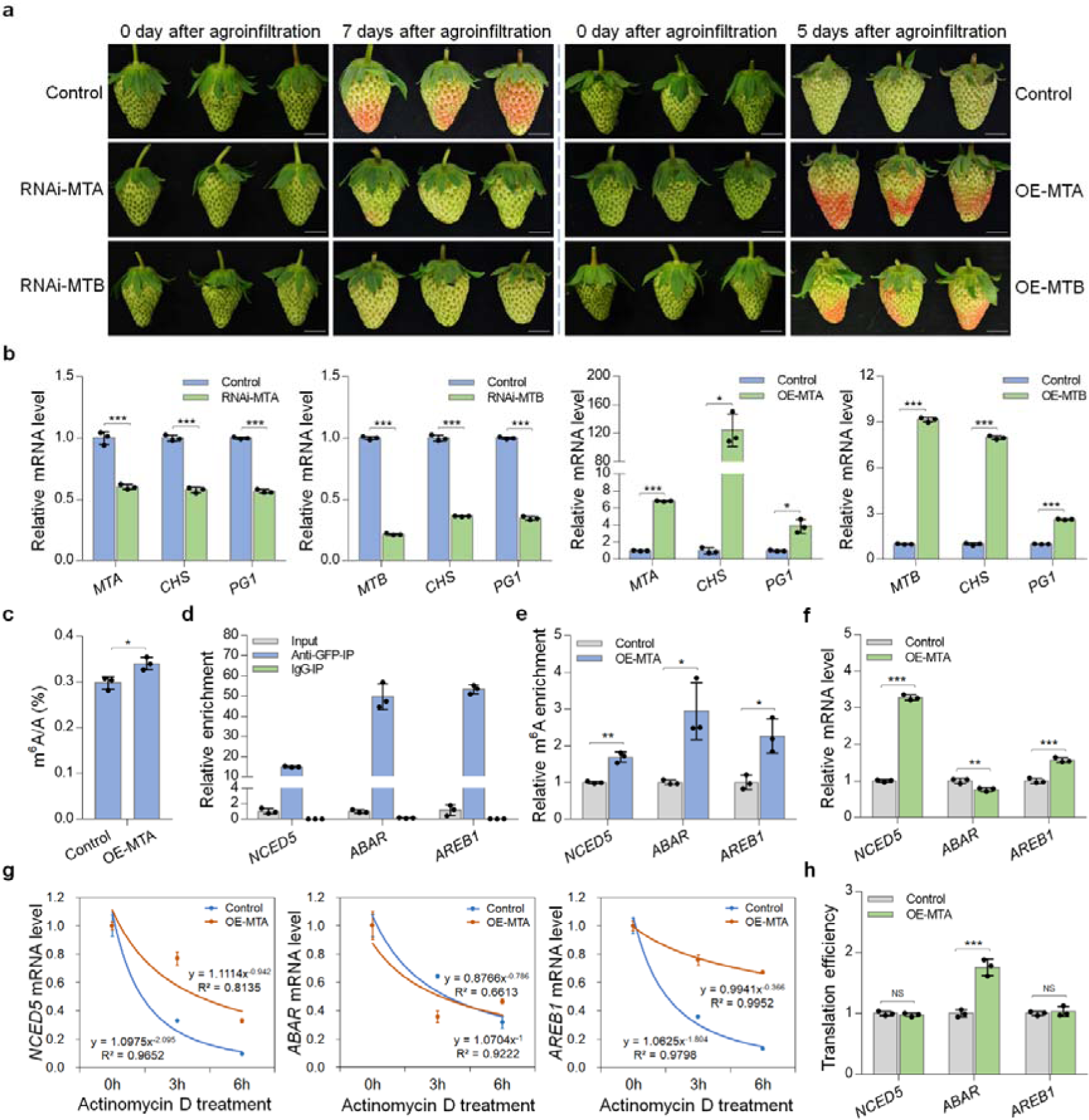
m^6^A methyltransferases positively regulate the ripening of strawberry fruit. **a** Ripening phenotypes of MTA/MTB RNA interference (RNAi-MTA/RNAi-MTB) and overexpression (OE-MTA/OE-MTB) fruits. Strawberry fruit agroinfiltrated with empty plasmid were used as controls. The representative images are shown. Scale bar = 1 cm. **b** Transcript levels of *MTA* and *MTB*, as well as the two important ripening genes *CHS* and *PG1*, in the RNAi and overexpression fruits determined by quantitative RT-PCR. The *ACTIN* gene served as an internal control. **c** LC-MS/MS assay revealing the changes in global m^6^A methylation levels in the MTA-overexpressed fruits. **d** RNA immunoprecipitation (RIP) assay revealing the binding of MTA protein to the transcripts of *NCED5*, *ABAR*, and *AREB1*. The protein-RNA complexes were extracted from strawberry fruit expressing the MTA-eGFP fusion protein and subjected to immunoprecipitation with anti-GFP monoclonal antibody or mouse IgG (negative control). **e**-**h** The changes in relative m^6^A enrichment (**e**), gene expression (**f**), mRNA stability (**g**), and translation efficiency (**h**) of *NCED5*, *ABAR*, and *AREB1* in the MTA-overexpressed fruits. The relative m^6^A enrichment and gene expression were determined by m^6^A-IP-qPCR and quantitative RT-PCR, respectively. For mRNA stability assay, the total RNAs were extracted after actinomycin D treatment at an indicated time point and submitted to quantitative RT-PCR assay. Translation efficiency was expressed as the abundance ratio of mRNA in the polysomal RNA versus the total RNA. Data are presented as mean ± standard deviation (n = 3). Asterisks indicate significant differences (\**P* < 0.05, \*\**P* < 0.01, \*\*\**P* < 0.001; Student’s t test). NS, no significance.

We next explored how the ripening of strawberry was regulated by MTA, the core component of the methyltransferase complexes with m^6^A catalytic activity. The *MTA*-overexpressed fruits displayed higher m^6^A levels than the control as revealed by LC-MS/MS (Fig. 6c). RNA immunoprecipitation (RIP) analysis indicated that the MTA directly binds to the transcripts of the ABA biosynthesis or signaling genes *NCED5*, *ABAR*, and *AREB1* (Fig. 6d). Accordingly, the m^6^A enrichments in the transcripts of *NCED5*, *ABAR*, and *AREB1* (Fig. 6e) and the corresponding mRNA levels (Fig. 6f) were significantly increased in the overexpressed fruits compared to the control. The mRNA stability assay demonstrated that mRNA of *NCED5* and *AREB1*, but not that of *ABAR*, degraded more slowly in the *MTA*-overexpressed fruits compared to the control (Fig. 6g). By contrast, *MTA* overexpression markedly enhanced the translation efficiency of the *ABAR*mRNA, but showed no significant effects on that of *NCED5* and *AREB1* (Fig. 6h). Together, these results suggest that MTA may regulate strawberry fruit ripening by targeting genes in the ABA pathway, leading to the increase in mRNA stability or translation efficiency of these genes.

### MTA-mediated m^6^A methylation modulates genes encoding translation initiation factors and elongation factors

Due to the critical role of ABA in the regulation of fruit ripening of strawberry, we evaluated whether *MTA* overexpression affects translation efficiency of other gene in the ABA signaling pathway. We found that genes, such as *WRKY DNA-binding protein 40* (*WRKY40*), exhibited significantly enhanced translation efficiency when the MTA was overexpressed (Additional file 1: Figure S8). This could not be reasonably explained by m^6^A deposition because the transcripts of these genes are not m^6^A-modified according to our m^6^A-seq datasets. We speculate that MTA may regulate translation efficiency of numerous transcripts beyond direct m^6^A installation. As expected, *MTA* overexpression also elevated the translation efficiency of a number of transcripts of ripening-related genes without m^6^A modification, such as *PG1* relevant to firmness and *dihydroflavonol 4-reductase* (*DFR*) associated with anthocyanin biosynthesis (Additional file 1: Figure S8).

To investigated the possible mechanisms, we mined our m^6^A-seq and RNA-seq data and found that the transcripts of genes encoding translation initiation factors (*EIF2*, *EIF2B*, *EIF3A*, and *EIF3C*) and elongation factors (*EF1A*), which play pivotal roles in facilitating protein synthesis by promoting the initiation and elongation of mRNA translation, respectively, exhibited m^6^A hypermethylation in the CDS region upon ripening initiation of strawberry fruit (Fig. 7a, b), concomitant with an increase in the transcript levels of these genes (Fig. 7c). Similar results were observed in the ripening process of the octoploid strawberry fruit (Fig. 7d, e). RIP analysis showed direct interactions between MTA and the transcripts of *EIF2*, *EIF2B*, *EIF3A*, and *EIF3C*, and *EF1A* (Fig. 7f). More importantly, the m^6^A enrichment in these gene transcripts (Fig. 7g) and the mRNA abundance (Fig. 7h) increased noticeably when *MTA* was overexpressed. The mRNA degradation rates were obviously reduced in the *MTA*-overexpressed fruits compared to the control (Fig. 7i), implying that *MTA*-mediated m^6^A modification stabilizes mRNAs of these genes. We propose that, in addition to direct modulation of translation efficiency via m^6^A installation in the target transcripts, MTA may indirectly modulate translation efficiency via m^6^A-mediated regulation of the translation initiation factors or elongation factors.

**Figure 7.**
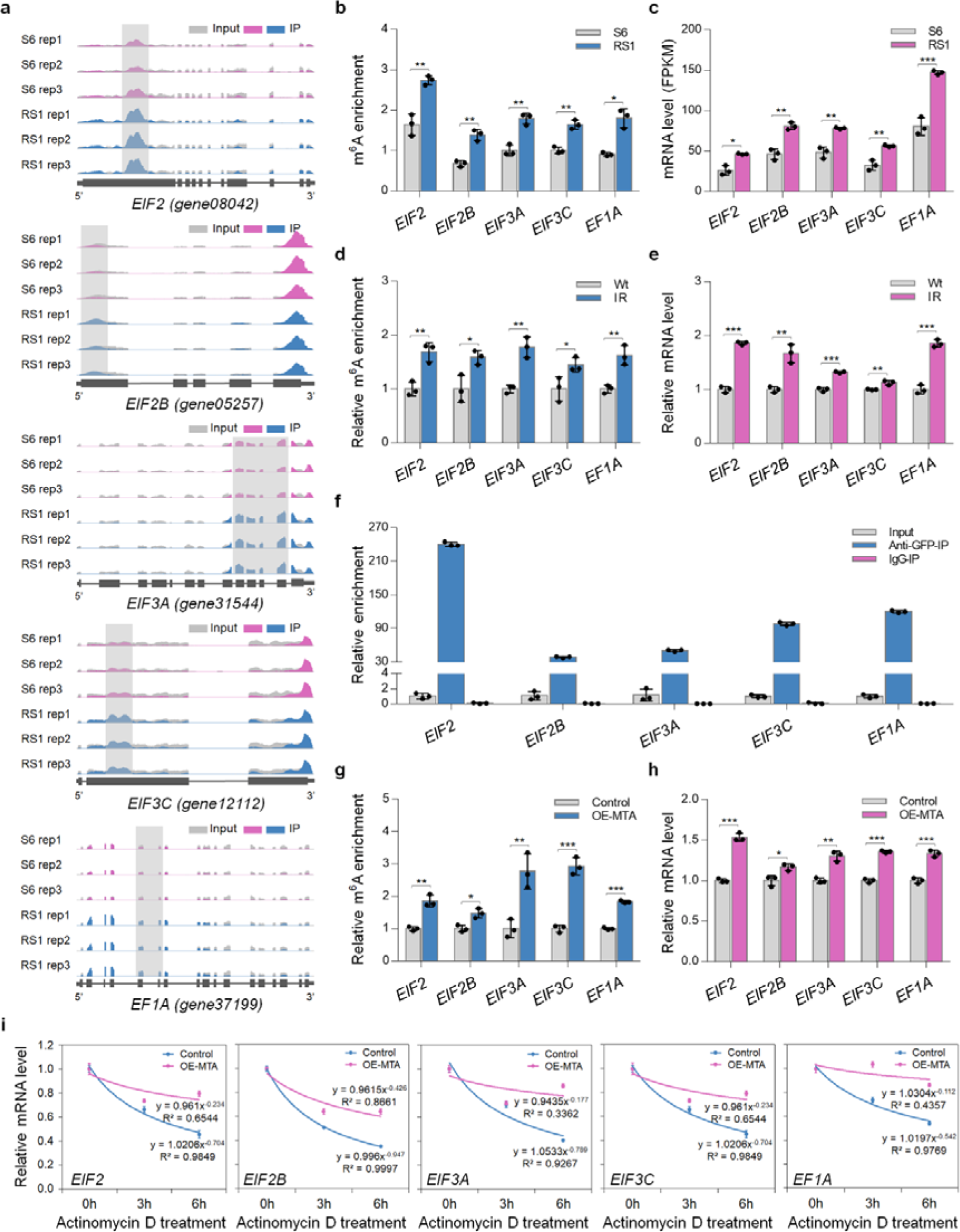
Influence of m^6^A on translation initiation factors or elongation factors in strawberry. **a** Integrated Genome Browser (IGB) tracks displaying the distribution of m^6^A reads in transcripts of genes encoding translation initiation factors (*EIF2*, *EIF2B*, *EIF3A*, and *EIF3C*) and elongation factors (*EF1A*). The hypermethylated m^6^A peaks (fold change ≥ 1.5; *P* value < 0.05) in fruit at RS1 stage compared to those at S6 stage are indicated by shadow boxes. S6, the growth stage 6; RS1, the ripening stage 1. Rep, replicate. **b** m^6^A enrichment for *EIF2*, *EIF2B*, *EIF3A*, *EIF3C*, and *EF1A* from m^6^A-seq data. **c** Transcript levels of *EIF2*, *EIF2B*, *EIF3A*, *EIF3C*, and *EF1A* determined by RNA-seq. FPKM, fragments per kilobase of exon per million mapped fragments. **d** and **e** Relative m^6^A enrichment (**d**) and gene expression (**e**) for *EIF2*, *EIF2B*, *EIF3A*, *EIF3C*, and *EF1A* in the octoploid strawberry fruit at the white (Wt) and initial red (IR) stages. The relative m^6^A enrichment and gene expression were determined by m^6^A-IP-qPCR and quantitative RT-PCR, respectively. The *ACTIN* gene served as an internal control. **f** RNA immunoprecipitation (RIP) assay revealing the binding of MTA protein to the transcripts of *EIF2*, *EIF2B*, *EIF3A*, *EIF3C*, and *EF1A*. **g**-**i** The changes in relative m^6^A enrichment (**g**), gene expression (**h**), and mRNA stability (**i**), of *EIF2*, *EIF2B*, *EIF3A*, *EIF3C*, and *EF1A* in the MTA-overexpressed fruits. For mRNA stability assay, the total RNAs were extracted after actinomycin D treatment at an indicated time point and subjected to quantitative RT-PCR assay. Data are presented as mean ± standard deviation (n = 3). Asterisks indicate significant differences (\**P* < 0.05, \*\**P* < 0.01, \*\*\**P* < 0.001; Student’s t test).

## Discussion

### m^6^A methylation participates in the regulation of strawberry fruit ripening

Compared to the understanding of the molecular basis underlying fruit ripening in climacteric fruits such as tomato, our knowledge regarding the regulation of ripening in non-climacteric fruits, e.g. strawberry, is still limited. Recently, it was elucidated that the ripening of strawberry involves the remodeling of DNA methylation [54]. However, it is unclear whether m^6^A methylation, which has been revealed to modulate the ripening of tomato fruit [36], is involved in the regulation of strawberry fruit ripening. In the present study, we found that m^6^A methylation represents a widespread mRNA modification in strawberry fruit and exhibits a dramatic change at the initiation stage of ripening. Overexpression of *MTA* or *MAB*, the m^6^A methyltransferase genes, promotes fruit ripening, whereas repression of either gene delays ripening (Fig. 6a, b), demonstrating that m^6^A methylation participates in the regulation of strawberry fruit ripening.

The transcript levels of *MTA* and *MTB* increased significantly upon ripening initiation of strawberry fruit (Fig. 5c-e), which may account for the m^6^A hypermethylation in the CDS region. It should be noted that the m^6^A methyltransferase genes in tomato express stably and no obviously global m^6^A hypermethylation was observed during fruit ripening [36]. By contrast, it is the m^6^A demethylase SlALKBH2 that positively regulates tomato fruit ripening through mediating the mRNA stability of *SlDML2*, a key DNA demethylase gene determining the DNA methylation patterns during ripening [36]. Due to the pivotal role of DNA methylation in the regulation of fruit ripening in both climacteric and non-climacteric fruits, it is reasonable to assume that m^6^A modification may modulate strawberry fruit ripening by modulating the DNA methylation machinery. Strawberry undergoes an overall loss of DNA methylation during ripening, and the reprogramming is governed by components in the RNA-directed DNA methylation (RdDM) pathway rather than those in the demethylation pathway [54]. However, our m^6^A-seq data indicated that there was no differential m^6^A modification in the transcripts of DNA methyltransferase genes in the RdDM pathway during strawberry fruit ripening (Additional file 1: Figure S9), suggesting that distinct mechanisms underlie the m^6^A-mediated ripening regulation in strawberry fruit.

### m^6^A methylation regulates strawberry fruit ripening by targeting ABA pathway

The phytohormone ABA plays crucial roles in plant growth, development, and stress responses [55]. The ripening of non-climacteric fruits, such as strawberry, has been revealed to be ABA-dependent [34]. Identification and functional definition of the core elements in ABA biosynthesis and signal transduction pathway have expanded our understanding of the mechanistic roles of ABA-mediated regulation of non-climacteric fruit ripening.

The ABA biosynthesis rate-limiting enzyme NCED was reported to play an essential role in the ripening of strawberry fruit [34, 38]. Moreover, several critical constituents of ABA signaling, including the ABA receptor FaPYR1 and FaABAR, the type 2C protein phosphatase FaABI1, and the SNF1-related kinase FaSnRK2.6, have been revealed to be indispensable for normal fruit ripening of strawberry [56–58]. Nevertheless, the regulatory mechanisms underlying ABA biosynthesis and signaling pathway remain largely unknown.

In *Arabidopsis*, ABA receptor PYLs could be regulated by post-translational modification, such as phosphorylation [59], tyrosine nitration [60], and ubiquitination [61], which synergistically modulate the abundance and activity of the receptors [55]. In contrast, the downstream signal molecules PP2Cs and SnRK2s are mainly controlled by protein phosphorylation and dephosphorylation for maintaining the appropriate ABA signal transduction [62–65]. The transcription of genes in ABA biosynthesis and signaling pathway displays dynamic changes in various development processes or in response to environmental stresses [55, 66–68], implying that they undergo precise regulation at transcriptional level. However, little is known about the regulation of genes in ABA pathway at the post-transcriptional level.

In this study, we found that *NCED5*, *ABAR*, and *AREB1*, the genes in ABA biosynthesis and signaling pathway undergo m^6^A-mediated post-transcriptional regulation (Fig. 4). The m^6^A modifications promote the mRNA stability of *NCED5* and *AREB1*, while enhance the translation efficiency of *ABAR*. These findings identify a novel layer of gene regulation in ABA biosynthesis and signaling pathway and establish a link between m^6^A-mediated ABA pathway and strawberry fruit ripening. Given the essential roles of ABA in plant development and stress resistance, it is interesting and necessary to explore the regulation of m^6^A methylation on these physiological processes.

### m^6^A modification exhibits diverse effects on mRNAs in strawberry

As the most prominent modification in mRNA, m^6^A was initially thought to accelerate degradation of mRNAs through promoting their transfer from the translatable pool to the decay sites [1]. Later findings have indicated that m^6^A modification also harbors the capacity to stabilize the mRNAs or promotes the translation in various biological processes [3, 22, 69, 70], although the underlying mechanisms are poorly understood, suggesting that m^6^A methylation possesses different regulatory roles on mRNAs [4]. This functional diversity was proposed to be heavily dominated by m^6^A distribution and the local sequence contexts within transcripts. One likely explanation is that the specific sequence contexts around m^6^A marks determine the recruitment of diverse m^6^A readers or other RNA binding proteins (RBPs) that carry distinguishing and even opposite molecular functions [4].

In this study, we observed a dramatic change in m^6^A modification at the initiation of strawberry fruit ripening (Fig. 2). The m^6^A depositions in the CDS region tend to stabilize the mRNAs, whereas those in the 3’ UTR or around the stop codon exhibit the opposite effects (Fig. 3). These data suggest that, depending on their distribution, the m^6^A modification plays distinct roles on mRNAs in strawberry. Notably, the differential m^6^A modification in the ripening process of tomato fruit mainly appeared around the stop codon or within the 3’ UTR, and these m^6^A depositions were generally negatively correlated with the abundance of the mRNAs. Combining the observation in strawberry and tomato, we propose that m^6^A modification around the stop codon or within the 3’ UTR tends to negatively correlated with mRNA abundance in the ripening process of both climacteric and non-climacteric fruits, whereas m^6^A deposition in the CDS region generally stabilize the mRNAs upon ripening initiation of non-climacteric fruits. Interestingly, besides modulating mRNA stability, m^6^A modification also affected translation efficiency of some transcripts. The changes in translation efficiency might be directly regulated by m^6^A deposition on the transcripts, or indirectly by m^6^A-mediated regulation of translation initiation factors and elongation factors.

In conclusion, our work reveals a regulatory role of m^6^A methylation on the ripening of the non-climacteric strawberry fruit. The molecular basis, which involves the m^6^A-mediated regulation of genes in the ABA biosynthesis and signaling pathway, is distinct from what was described in climacteric tomato fruit (Additional file 1: Figure S10). These findings provide new insights into understanding the regulatory networks controlling fruit ripening.

## Methods

### Plant materials

The diploid woodland strawberry (*Fragaria vesca* cv. ‘Hawaii-4’) and the octoploid cultivated strawberry (*Fragaria × ananassa* cv. ‘Benihoppe’) were planted in a greenhouse with standard culture conditions [58]. To accurately determine fruit ages through development, flowers were tagged at the anthesis. Fruits of ‘Hawaii-4’ at the growth stage 6 (S6), the ripening stage 1 (RS1), and the ripening stage 3 (RS3) [38], which were on average 15, 21, and 27 days post-anthesis (DPA), respectively, were harvested, and then frozen in liquid nitrogen. The fruits with the removal of attached achenes were subsequently stored at −80 °C until use. The ‘Benihoppe’ fruits were harvested at the small green (SG), large green (LG), white (Wt), initial red (IR), and full red (FR) stages, respectively, based on the size, weight, shape, and color [71], and then maintained at fresh status for further studies or directly frozen and stored as the ‘Hawaii-4’.

### m^6^A-seq and data analysis

The m^6^A-seq was performed according to the method described by Dominissini et al (2013) [37]. Briefly, total RNAs were extracted from the woodland strawberry fruit at S6, RS1, and RS3 stages by the plant RNA extraction kit (Magen, R4165-02), and then 300 µg of intact total RNAs were used for mRNA isolation by the Dynabeads mRNA purification kit (Life Technologies, 61006). The purified mRNAs were randomly fragmented into ∼100 nucleotide-long fragments by incubation at 94 °C for 5 min in the RNA fragmentation buffer (10 mM Tris-HCl, pH 7.0, and 10 mM ZnCl_2_). The reaction was terminated with 50 mM EDTA, and then the fragmented mRNAs were purified by standard phenol-chloroform extraction and ethanol precipitation. For immunoprecipitation (IP), 5 µg of fragmented mRNAs was mixed with 10 µg of anti-m^6^A polyclonal antibody (Synaptic Systems, 202003) and incubated at 4 °C for 2 h in 450 µ L of IP buffer consisting of 10 mM Tris-HCl, pH 7.4, 150 mM NaCl, 0.1% NP-40 (v/v), and 300 U mL^-1^ RNase inhibitor (Promega, N2112S). After the addition of 50 µL Dynabeads protein-A (Life Technologies, 10002A), the mixture was incubated at 4 °C for another 2 h. The beads were subsequently washed twice with high-salt buffer containing 50 mM Tris-HCl, pH 7.4, 1 M NaCl, 1 mM EDTA, 1% NP-40 (v/v), and 0.1% SDS (w/v) and twice with IP buffer. The m^6^A-containing fragments were eluted from the beads by incubation with 6.7 mM N^6^-methyladenosine (TargetMol, T6599) in IP buffer at 4 °C for 2 h, followed by standard phenol-chloroform extraction and ethanol precipitation. Then, 50 ng of m^6^A-containing mRNAs or pre-immunoprecipitated mRNAs (the input) were used for library construction by the NEBNext ultra RNA library preparation kit (NEB, E7530). High-throughput sequencing was conducted on the illumina HiSeq X sequencer with a paired-end read length of 150 bp following the standard protocols. The sequencing was performed with three independent biological replicates, and each RNA sample was prepared from the mix of at least 60 strawberry fruits to avoid individual difference among fruits.

For data analysis, the quality of raw sequencing reads was initially assessed by FastQC tool (version 0.11.7; http://www.bioinformatics.babraham.ac.uk/projects/fastqc). Adaptors and low-quality bases with a score < 20 were trimmed using Cutadapt (version 1.16) [72], and then short reads with a length < 18 nucleotides or reads containing ambiguous nucleotides were filtered out by Trimmomatic (version 0.30) [73]. The remaining reads were aligned to the strawberry reference genome v1.1 (ftp://ftp.bioinfo.wsu.edu/species/Fragaria_vesca/Fvesca-genome.v1.1) by Burrows Wheeler Aligner (BWA; version 0.30) [74]. Mapping quality (MAPQ) of all aligned reads was concurrently evaluated, and only uniquely mapped reads with a MAPQ ≥ 13 were retained for subsequent analysis [37].

The identification of m^6^A peaks was carried out by MACS software (version 2.0.10) [75], using the corresponding input as a control. High-confidence peaks were obtained by a stringent cutoff threshold for MACS-assigned false discovery rate (FDR) < 0.05. PeakAnalyzer (version 2.0) [76] was applied to annotate the identified peaks to the strawberry genome annotation file (ftp.bioinfo.wsu.edu/species/Fragaria_vesca/Fvesca-genome.v2.0.a2/genes/). To identify the differentially methylated peaks between samples, the m^6^A site differential algorithm [77] was applied with a criterion of fold change in m^6^A enrichment ≥ 1.5 and *P* value < 0.05. HOMER (version 4.7; http://homer.ucsd.edu/homer/) [78] was employed to identify the m^6^A motifs with a restricted length of six nucleotides. The differential m^6^A peaks identified between S6 and RS1 stages were used as the target sequences, and the exon sequences without m^6^A peaks were used as the background sequences. Integrated Genome Browser (IGB, version 9.0.2) [79] was used for visualization of m^6^A peaks. Gene Ontology (GO) enrichment was analyzed on the agriGO database (version 2.0; http://systemsbiology.cau.edu.cn/agriGOv2/) and only statistically significant terms with a Yekutieli-corrected *P* value < 0.05 were remained [80].

### RNA-seq and data analysis

The sequencing reads from the input samples in m^6^A-seq were used for RNA-seq analysis as previously reported [21]. In brief, the uniquely mapped reads with a MAPQ ≥ 13 were assembled by Cufflinks [81]. Gene expression was presented as fragments per kilobase of exon per million mapped fragments (FPKM) by using Cuffdiff [81], which concurrently provides statistical routines for capturing differentially expressed genes. The Benjamini and Hochberg’s approach [82] was used to adjust the resulting *P* values for controlling the false discovery rate (FDR). Differential gene expression was defined on basis of a cutoff criterion of FPKM fold change ≥ 1.5 and *P* value < 0.05.

### Quantitative RT-PCR analysis

Total RNAs were extracted from the strawberry fruits or *N. benthamiana* leaves using the plant RNA extraction kit (Magen, R4165-02). The extracted RNAs were reverse transcribed into cDNAs by the HiScript^®^ III RT SuperMix for qPCR kit (Vazyme, R323-01). The synthesized cDNAs were then employed as templates for PCR amplification using the ChemQ Universal SYBR qPCR Master Mix (Vazyme, Q711-02-AA) and a StepOne Plus Real-Time PCR System (Applied Biosystems) in the following program: 95 °C for 10 min, followed by 40 cycles of 95 °C for 15 s and 60 °C for 30 s. Relative quantification of gene transcription levels were performed by the cycle threshold (C_T_)2^(-ΔCT)^ method [83].

Strawberry *ACTIN* (gene22626) or *N. benthamiana ACTIN* (*Niben101Scf03410g03002*) was applied to normalize the expression values. The primers for PCR amplification are listed in Additional file 2: Table S17. The experiment was conducted with three biological replicates, and each contained three technical repeats.

### m^6^A-IP-qPCR

m^6^A-IP-qPCR was carried out as previously described with minor modifications [84]. Briefly, 5 µg of intact mRNAs were fragmented into ∼300 nucleotide-long fragments by an incubation at 94 °C for 30 s in the RNA fragmentation buffer (10 mM Tris-HCl, pH 7.0, and 10 mM ZnCl_2_), followed by the addition of 50 mM EDTA to terminate the reaction. The fragmented mRNAs were purified by standard ethanol precipitation and resuspended in 250 μL of DEPC-treated water. Then, 5 μL of fragmented mRNAs was taken as the input control and 100 μL were incubated with 5 µg of anti-m^6^A polyclonal antibody (Synaptic Systems, 202003) at 4 °C for 2 h in 450 µL of IP buffer containing 10 mM Tris-HCl, pH 7.4, 150 mM NaCl, 0.1% NP-40 (v/v), and 300 U mL^-1^ RNase inhibitor (Promega, N2112S). The mixture was subsequently incubated with 20 µL of Dynabeads protein-A (Life Technologies, 10002A) at 4 °C for another 2 h. After washing twice with high-salt buffer containing 50 mM Tris-HCl, pH 7.4, 1 M NaCl, 1 mM EDTA, 1% NP-40 (v/v), and 0.1% SDS (w/v) and twice with IP buffer, the m^6^A-containing fragments were eluted by an incubation with 6.7 mM N^6^-methyladenosine (m^6^A; TargetMol, T6599) in 200 µL of IP buffer at 4 °C for 2 h, followed by ethanol precipitation. The immunoprecipitated mRNA fragments were finally resuspended in 5 μL DEPC-treated water. Then, both the m^6^A-containing mRNAs and the input mRNAs were submitted to quantitative RT-PCR by using the primers listed in Additional file 2: Table S17. Relative m^6^A enrichment in specific region of a transcript was calculated using the cycle threshold (C_T_) 2^(−ΔCT)^ method [83]. The value for the immunoprecipitated sample was normalized against that for the input. The experiment was performed with three biological replicates, and each contained three technical repeats.

### Identification of strawberry m^6^A methyltransferases and phylogenetic analysis

To identify m^6^A methyltransferases in strawberry, the amino acid sequence of conserved MT-A70 domain (PF05063) was downloaded from the Pfam (http://pfam.xfam.org/) and then utilized to search the potential homologs against the strawberry protein dataset (ftp.bioinfo.wsu.edu/species/Fragaria_vesca/Fvesca-genome.v2.0.a2/genes/) using HMMER 3.1 with default parameters [85]. The resulting protein sequences were analyzed on the CDD database (https://www.ncbi.nlm.nih.gov/cdd/) [86], and only that containing a MT-A70 domain were remained as the m^6^A methyltransferase candidates. The known m^6^A methyltransferases METTL3 and METTL14 in mammals and MTA and MTB in *Arabidopsis* were then employed to perform BLASTP-algorithm in NCBI (https://www.ncbi.nlm.nih.gov/) to further confirm the identified homologs. For phylogenetic analysis, the sequences of the identified strawberry m^6^A methyltransferase were aligned with the sequences of m^6^A methyltransferase in mouse (*Mus musculus*), rice (*Oryza sativa*), maize (*Zea mays*), tomato (*Solanum lycopersicum*), tobacco (*Nicotiana benthamiana*), and *Arabidopsis*, using ClustalX 2.1 with standard parameters [87]. The alignment result was imported into MEGA software (version 5.2) to create the phylogenetic tree by using Neighbor-Joining method with 1000 bootstrap replicates [88].

### Y2H analysis

Y2H analysis was performed as the previously described [89]. In brief, the coding sequence of *MTA* and *MTB* was amplified from the cDNAs of diploid woodland strawberry and then ligated into the pGADT7 (AD) and pGBKT7 (BD) vectors, respectively. The resulting plasmids were co-transformed into *Saccharomyces cerevisiae* strain AH109 according to the protocols in the Matchmaker GAL4 Two-Hybrid System 3 (Clontech). The yeast cells were cultured on SD/-Leu-Trp (-LW) medium, and then transferred onto the SD/-Leu-Trp-His (-LWH) or SD/-Leu-Trp-His-Ade medium (-LWHA) with or without X-α-gal. Transformants carrying empty pGADT7 (AD) or pGBKT7 (BD) vectors were used as negative controls, and that concurrently carrying pGBKT7-53 and pGADT7-T vectors was used as the positive control. The primers used for vector constructions are listed in Additional file 2: Table S17.

### LCI assay

LCI assay was performed as previously reported [90]. Briefly, the coding sequence of *MTA* was cloned into the pCambia1300-nLUC plasmid, and the coding sequence of *MTB* or its MT-A70 domain sequence (*MTB^D^*) was cloned into the pCambia1300-cLUC plasmid. The resulting constructs were separately transformed into *Agrobacterium tumefaciens* strain GV3101. The agrobacteria were cultured at 28 °C for 18 h in Luria-Bertani (LB) liquid medium containing 50 μg mL^−1^ kanamycin, 50 μg mL^−^ gentamycin, and 50 μg mL^−1^ rifampicin. After being pelleted by centrifugation at 5,000 g for 5 min, the agrobacteria were resuspended in the infiltration buffer (10 mM MES, pH 5.6, 10 mM MgCl_2_, and 100 µM acetosyringone) to a final OD_600_ of 0.5. Then, the suspension was infiltrated into *N. benthamiana* leaves for co-expression of fusion protein nLUC-MTA with MTB-cLUC or MTB^D^-cLUC. After culture for 36 h, the leaves were incubated with 1 mM luciferin dissolved in ddH_2_O supplemented with 0.01% Triton X-100 at room temperature for 5min, and then observed under a chemiluminescence imaging system (Tanon). Empty vectors expressing cLUC or nLUC were co-transformed as the negative controls. The primers used for vector constructions are listed in Additional file 2: Table S17.

### Subcellular localization

For subcellular localization analysis, the coding sequence of *MTA* and *MTB* was amplified from the cDNAs of diploid woodland strawberry and then inserted into the pCambia2300-mCherry and pCambia2300-eGFP plasmids to generate 35S::*MTA-mCherry* and 35S::*MTB-eGFP* vectors, respectively. The resulting constructs were separately transformed into *A. tumefaciens* strain GV3101. The agrobacteria were subsequently infiltrated into *N. benthamiana* leaves for the individual expression of mCherry-tagged MTA (MTA-mCherry) and eGFP-tagged MTB (MTB-eGFP) or the co-expression of the two fusion proteins. After culture for 36 h, the mesophyll protoplasts were isolated from *N. benthamiana* leaves as previously reported [91] and observed under a Leica confocal microscope (Leica DMI600CS). Protoplasts expressing eGFP or mCherry were used as negative controls. The primers used for vector constructions are listed in Additional file 2: Table S17.

### Agroinfiltration-mediated transient transformation in strawberry fruit

Transient transformation of strawberry fruit mediated by agroinfiltration was performed as previously described [53]. To construct the RNA interference (RNAi) vectors, a ∼300 bp fragment targeting the coding sequence region of *MTA* or *MTB* was cloned and inserted into the pCR8 plasmid, and then restructured into the pK7GWIWGD (II) plasmid by using the Gateway LR Clonase^TM^ Enzyme Mix (Invitrogen, 11791-020). To construct the overexpression (OE) vectors, the coding sequence of *MTA* and *MTB* was amplified and ligated into the pCambia2300-eGFP plasmid to generate 35S::*MTA-eGFP* and 35S::*MTB-eGFP* vector, respectively. The resulting constructs were separately transformed into the *A. tumefaciens* strain GV3101. The agrobacteria were cultured at 28 °C overnight in LB liquid medium supplemented with 50 μg mL^−1^ kanamycin, 50 μg mL^−1^ gentamycin, and 50 μg mL^−1^ rifampicin, and then diluted 1:100 in 100 mL of fresh LB medium to continue culturing for approximately 8 h. The agrobacteria cells were subsequently collected by centrifugation at 5,000 g for 5 min and resuspended in the infiltration buffer (10 mM MES, pH 5.6, 10 mM MgCl_2_, and 100 µM acetosyringone) to a final OD_600_ of 0.8. After being kept at room temperature for 2 h without shaking, the suspensions were injected into the octoploid strawberry fruit at large green (LG) stage by using a 1 mL syringe. The infiltrated fruits were cultured for 5-7 days in a growth room with the following conditions: 23 °C, 80 % relative humidity, and a 16/8 h light/dark photoperiod with a light intensity of 100 μmol ms^-1^. The experiment was performed with three independent biological replicates, and each group contained at least fifteen fruits. The primers used for vector constructions are listed in Additional file 2: Table S17.

### mRNA stability assay

For mRNA stability assay in *N. benthamiana* leaves, the coding sequence of *NCED5*, *ABAR*, and *AREB1* was amplified from the cDNAs of diploid woodland strawberry. The mutated forms of the amplified sequence in which the potential m^6^A sites were replaced by cytidine (C) or guanine (G) were constructed using the QuikChange II XL Site-directed Mutagenesis Kit (Agilent Technologies, 200518). The fragments were separately inserted into the pCambia2300 vector, which were subsequently transformed into *A. tumefaciens* strain GV3101. Then, the agrobacteria were cultured at 28 °C for 18 h in LB liquid medium containing 50 μg mL^−1^ kanamycin, 50 μg mL^−1^ gentamycin, and 50 μg mL^−1^ rifampicin. After being collected by centrifugation at 5,000 g for 5 min, the agrobacteria were diluted to an OD_600_ of 0.5 in the infiltration buffer (10 mM MES, pH 5.6, 10 mM MgCl_2_, 100 µM acetosyringone), and then infiltrated into the *N. benthamiana* leaves. After 36 h of incubation, the infiltration parts in the leaves were injected with 20 µg mL^−1^ actinomycin D (Sigma, A4262) dissolved in ddH_2_O. After culture for 30 min, leaf discs were taken and considered as time 0 controls, and subsequent samples were harvested every 3 h in triplicate.

For mRNA stability assay in strawberry, the *MTA*-overexpressed fruit and the control were sliced into ∼3 mm slices, and then transferred onto plates containing 20 μg mL^−1^ actinomycin D dissolved in ddH_2_O. After incubation for 30 min, the slices were collected as the *N. benthamiana* leaf discs. The mRNA levels of genes were subsequently examined by quantitative RT-PCR as described above. All primers used for PCR amplification were listed in Additional file 2: Table S17.

### Translation efficiency assay

Translation efficiency was assayed according to the method described by Merchante et al (2015) [49]. Briefly, 6 g of strawberry fruits or 3 g of *N. benthamiana* leaves were ground into fine powder in liquid nitrogen. One gram of sample was used for total RNA extraction, and the rest was suspended in 15 mL of polysome extraction buffer (200 mM Tris-HCl, pH 9.0, 35 mM MgCl_2_, 200 mM KCl, 25 mM EGTA, 1% Triton X-100 (v/v), 1% IGEPAL CA-630 (v/v), 5 mM DTT, 1 mM PMSF, 50 μg mL^-1^ Chloramphenicol, and 100 μg mL^-1^ Cycloheximide) at 4 °C for 20 min with slight shaking. The mixture was centrifuged twice at 16, 000 g for 20 min at 4 °C. Then, 12.5 mL of supernatant were slowly transferred onto 13.5 mL of sucrose buffer (1.75 M Sucrose, 400 mM Tris-HCl, pH 9.0, 35 mM MgCl_2_, 5 mM EGTA, 200 mM KCl, 5 mM DTT, 50 μg mL^-1^ Chloramphenicol, and 50 μg ml^-1^ Cycloheximide). After centrifugation at 200,000 g for 4 h at 4 °C, the supernatant was carefully removed, and the polysomes in the bottom were resuspended in 300 μL of DEPC-treated water. The polysomal RNAs, as well as the total RNAs, were isolated by using the plant RNA extraction kit (Magen, R4165-02), and used for quantitative RT-PCR analysis as described above. Translation efficiency was calculated by the abundance ratio of mRNA in the polysomal RNA versus the total RNA using the cycle threshold (CT) 2^(− CT)^ method [83] with the *GADPH2* gene as an internal reference. The primers used for PCR amplification are listed in Additional file 2: Table S17.

### Quantitative analysis of m^6^A level by LC-MS/MS

The global m^6^A levels in strawberry fruit were detected by LC-MS/MS as previously described [36] with minor modifications. In brief, 200 ng of mRNAs was digested at 37 °C for 6 h with 1 unit of Nuclease P1 (Wako, 145-08221) in 50 μL of reaction buffer containing 10 mM ammonium acetate, pH 5.3, 25 mM NaCl, and 2.5 mM ZnCl_2_. Then, 1 unit of alkaline phosphatase (Sigma-Aldrich, P6774) and 5.5 μL of 1 M fresh NH_4_HCO_3_ were added, followed by incubation at 37 °C for another 6 h. After centrifugation at 15,000 g for 5 min, the supernatant was used for LC-MS/MS analysis. The digested nucleosides were separated by UPLC (Waters, ACQUITY) equipped with a ACQUITY UPLC HSS T3 column (Waters), and then detected by a Triple Quad Xevo TQ-S (Waters) mass spectrometer in positive ion mode with multiple reaction monitoring. The mobile phase was composed of buffer A (0.1% formic acid in ultrapure water) and buffer B (100% acetonitrile). Nucleosides were accurately quantified depending on the nucleoside-to-base ion mass transitions of m/z 268.0 to 136.0 (A) and m/z 282.0 to 150.1 (m^6^A). The pure commercial adenosine (A; TargetMol, T0853) and N^6^-methyladenosine (m^6^A; TargetMol, T6599) were used to generate standard curves, which were subsequently employed to calculate the contents of A and m^6^A in each sample. The global m^6^A levels were presented in the form of m^6^A/A ratio. The experiment was repeated with three independent biological replicates.

### RNA immunoprecipitation

RNA immunoprecipitation (RIP) was carried out following the method of Wei et al (2018) [22] with minor modifications. Briefly, the octoploid strawberry fruit expressing the MTA-eGFP protein were sliced into ∼2 mm slices, and then fixed with 1% formaldehyde under a vacuum for 30 min on ice. The fixation was terminated by the addition of 150 mM glycine, followed by incubation on ice for 5 min. The fixed fruit tissues (2 g) were homogenized in 5 mL of lysis buffer (50 mM HEPES, pH 7.5, 2 mM EDTA, 150 mM KCl, 0.5% NP-40 (v/v), 2 mM EDTA, 0.5 mM DTT, 1× cocktail protease inhibitor (Sigma, 04693132001), and 300 U mL ^−1^ RNase Inhibitor). After incubation at 4 °C for 1 h, the mixture was centrifuged at 15,000 g for 30 min. Then, 200 μL of the supernatant were taken as the input control, and the remainder was subjected to immunoprecipitation (IP) with anti-GFP monoclonal antibody or rabbit lgG at 4 °C overnight. Fifty microliters of Dynabeads Protein-A (Life Technologies, 10002A) were added to the mixture and incubated at 4 °C for 2 h. After washing four times with PBS buffer, the RNA-protein mix was catalyzed by proteinase K (Takara, 9034) at 55 °C for 1 h. The immunoprecipitated mRNAs and input mRNAs were subsequently isolated by the plant RNA extraction kit (Magen, R4165-02). Relative enrichment of individual transcript was determined by quantitative RT-PCR analysis as described above. The primers used for PCR amplification are listed in Additional file 2: Table S17. The analysis was performed with three biological replicates, and each contained three technical repeats.

### Data access

The raw sequencing data and processed peaks data in m^6^A-seq have been deposited in the Gene Expression Omnibus database under the accession number GSE167183. All the other data generated in this study are included in the article and the Additional files.

## Additional material

**Additional file 1:** Supplementary Figure S1-S10. **Figure S1**. Pearson correlation analysis of input reads in the m^6^A peak regions identified from m^6^A-seq. **Figure S2**. Pearson correlation analysis of immunoprecipitation (IP) reads in the m^6^A peak regions identified from m^6^A-seq. **Figure S3**. Validation of confident m^6^A peaks. **Figure S4**. Sequence motif identified within the m^6^A peaks by HOMER (http://homer.ucsd.edu/homer/). **Figure S5**. m^6^A modification and expression of genes in ABA biosynthesis and signaling pathway in octoploid cultivated strawberry. **Figure S6**. Identification of the conserved MT-A70 domain of m^6^A methyltransferases MTA and MTB in diploid woodland strawberry and octoploid cultivated strawberry. **Figure S7**. Protein sequence alignment of the MT-A70 domains in m^6^A methyltransferases. **Figure S8**. The changes in translation efficiency of ABA signaling gene and ripening genes in the MTA-overexpressed fruits. **Figure S9**. m^6^A enrichment for DNA methyltransferase genes from m^6^A-seq data. **Figure S10**. Regulatory model for the m6A-mediated ripening in climacteric tomato fruit and non-climacteric strawberry fruit.

**Additional file 2:** Supplementary Table S1-S17. **Table S1**. A summary of m^6^A-seq information in strawberry fruit at different stages. **Table S2**. Confident m^6^A peaks identified by three independent m^6^A-seq experiments in strawberry fruit at S6 stage. **Table S3**. Confident m^6^A peaks identified by three independent m^6^A-seq experiments in strawberry fruit at RS1 stage. **Table S4**. Confident m^6^A peaks identified by three independent m^6^A-seq experiments in strawberry fruit at RS3 stage. **Table S5**. m^6^A density in actively expressed transcripts. **Table S6**. Transcripts containing hypermethylated m^6^A peak in strawberry fruit at RS1 stage compared to those at S6 stage. **Table S7**. Transcripts containing hypomethylated m^6^A peak in strawberry fruit at RS1 stage compared to those at S6 stage. **Table S8**. Transcripts containing hypermethylated m^6^A peak in strawberry fruit at RS3 stage compared to those at RS1 fruit. **Table S9**. Transcripts containing hypomethylated m^6^A peak in strawberry fruit at RS3 stage compared to those at RS1 stage. **Table S10**. Ripening-specific peaks identified in strawberry fruit at RS1 stage. **Table S11**. Differentially expressed genes between strawberry fruit at S6 and RS1 stage identified by three independent RNA-seq experiments. **Table S12**. Differentially expressed genes between strawberry fruit at RS1 and RS3 stage identified by three independent RNA-seq experiments. **Table S13**. Differentially expressed genes containing hypermethylated m^6^A peaks in strawberry fruit at RS1 stage compared to those at S6 stage. **Table S14**. Differentially expressed genes containing hypomethylated m^6^A peaks in strawberry fruit at RS1 stage compared to those at S6 stage. **Table S15**. Differentially expressed genes containing hypermethylated m^6^A peaks in strawberry fruit at RS3 stage compared to those at RS1 stage. **Table S16**. Differentially expressed genes containing hypomethylated m^6^A peaks in strawberry fruit at RS3 stage compared to those at RS1 stage. **Table S17**. A summary of primer informations.

## Abbreviations

ABA: abscisic acid
*ABAR*: *putative ABA receptor*
AD: activation domain
ALKBH: AlkB homolog
*AREB1*: *ABA-responsive element-binding protein 1*
BD: binding domain
CDS: coding sequence
*CHS*: *chalcone synthase*
*DFR*: *dihydroflavonol 4-reductase*
DPA: days post-anthesis
EF: translation elongation factor
eGFP: enhanced green fluorescent protein
EIF: translation initiation factor
FDR: false discovery rate
FPKM: fragments per kilobase of transcript per million fragments mapped
FR: full red
FTO: fat mass and obesity-associated protein
GO: gene ontology
HOMER: hypergeometric optimization of motif enrichment
IGB: Integrated Genome Browser
IP: immunoprecipitation
IR: initial red
LB: Luria-Bertani
LCI: luciferase complementation imaging
LG: large green
m^6^A: N^6^-methyladenosine
MAPQ: mapping quality
METTL: methyltransferase like
mRNA: messenger RNA
MTA: adenosine methyltransferase
MTB: adenosine methyltransferase B
*NCED5*: *9-cis-epoxycarotenoid dioxygenase 5*
OE: overexpression
*PG1*: *polygalacturonase 1*
RBP: RNA binding proteins
RdDM: RNA-directed DNA methylation
RIP: RNA immunoprecipitation
RNAi: RNA interference
RS1: the ripening stage 1
RS3: the ripening stage 3
RT-PCR: reverse transcription polymerase chain reaction; S6 the growth stage 6
SG: small green
TSS: transcription start site
UTR: untranslated region
*WRKY40*: *WRKY DNA-binding protein 40*
Wt: white
WT: wild-type
WTAP: Wilms’ tumor 1-associating protein
Y2H: yeast two-hybrid.

## Acknowledgements

We thank Hang Su (Institute of Botany, Chinese Academy of Sciences) for the assistance with LC-MS/MS assay. We thank Jingquan Li (Institute of Botany, Chinese Academy of Sciences) for the assistance with confocal microscopy assay. We also thank Jianmin Zhou (Institute of Genetics and Developmental Biology, Chinese Academy of Sciences) for providing the pCambia1300–nLUC/cLUC vectors.

## Funding

This work was supported by the National Key Research and Development Program (2018YFD1000200) and the National Natural Science Foundation of China (grant Nos. 31925035, 31930086, and 31572174).

## Availability of data and materials

The raw sequencing data and processed peaks data in m^6^A-seq have been deposited in the Gene Expression Omnibus database under the accession number GSE167183 (https://www.ncbi.nlm.nih.gov/geo/query/acc.cgi?acc=GSE167183) with a temporary token of yjatoqksvribzkf. All the other data generated in this study are included in the article and the Additional files.

## Authors’ contributions

GQ conceived, designed, and supervised the experiments. ST and BL provided critical discussions. LZ and RT performed the experiments and analyzed the data. LX prepared the strawberry materials. GQ and LZ wrote the manuscript with contributions from RT. All authors read and approved the final manuscript.

## Competing interests

The authors declare that they have no competing interests.

## Consent for publication

Not applicable.

## Ethics approval and consent to participate

Not applicable.

## Notes

### Competing Interest Statement

The authors have declared no competing interest.

